# Arbitrium communication controls phage life-cycle through modulation of a bacterial anti-phage defense system

**DOI:** 10.1101/2023.04.27.537455

**Authors:** Polina Guler, Shira Omer Bendori, Nitzan Aframian, Amit Kessel, Avigdor Eldar

## Abstract

Bacterial temperate viruses (phages) have to decide between a quiescent (lysogenic) and virulent (lytic) lifestyle in the face of a variety of phage defense systems. Multiple *Bacilli* phage families have been shown to use the arbitrium communication system, but the mechanism by which the arbitrium system exerts its function remains largely unknown. Here we study phage ɸ3T, in which arbitrium was originally identified, and find that arbitrium communication controls the phage life-cycle through interactions with a host-encoded defense system. Under lytic conditions, the arbitrium system expresses an anti-toxin, AimX, which blocks the RNA ribonuclease activity of MazF, part of the MazEF toxin-antitoxin system. When arbitrium signal concentration is high, AimX is not expressed and MazF remains active. We find that this activity is necessary for lysogenization. Finally, we show that MazEF acts as a defense system, and protects bacteria against a lytic ɸ3T mutant which lacks AimX and an additional later-expressed MazE-like antitoxin, YosL. Altogether, our results show how a bacterial defense system has been co-opted by phages to control their lysis/lysogeny decision-making.

## Introduction

Temperate bacteriophages can transition between two major stages of their life cycle, a quiescent stage (lysogeny) where they typically integrate into the genome of their bacterial host, and a virulent stage (lysis), where they replicate within their host to form multiple virions which are released by lysing the host cell. Phages have evolved complex regulatory networks to control whether to become lytic or lysogenic upon initial infection (lysis/lysogeny) or when integrated as a prophage in the bacterial genome (prophage induction)^1,2^.

Recently, it was found that in many *Bacillus* phages both lysis/lysogeny and prophage induction decisions are regulated by a peptide-based cell-cell signaling system named arbitrium ^3–6^. This system, which belongs to the RRNPP super-family of quorum-sensing systems ^7,8^, was mostly studied in the SPβ-like phages ɸ3T and SPβ ^3–6,9–12^, but exists in many other phage families ^6,13^. The arbitrium system consists of the AimR receptor, the AimP peptide signal and the *aimX* lysis promoting gene. AimR and AimP are expressed both upon infection and in the lysogen. AimP pre-pro-peptide is secreted and cleaved twice to form the mature AimP peptide which can be imported back into the cell cytoplasm by oligopeptide permease systems. The AimR receptor promotes the expression of the *aimX* gene in the absence of peptide, while peptide binding prevents AimR activity ^3,9–12,14^. During infection, activation of the *aimX* gene is necessary to promote lysis while during prophage induction, *aimX* activity is integrated with a DNA damage signal to yield the final prophage induction decision ^3–6^. How *aimX* promotes lysis is currently unclear. In SPβ-like phages it was proposed that it functions as a non-coding RNA in a yet uncharacterized manner, as there is no clear open reading frame (ORF) in phage SPβ ^3^. In the related phage ɸ3T, there exists a small ORF within the *aimX* gene, but it is not known whether this putative resulting peptide is functionally important ^3^.

Phages face a variety of phage defense systems employed by their hosts ^15–20^. These include both systems which directly restrict phages such as restriction-modification and CRISPR systems as well as a large cohort of abortive infection systems. In the latter type, in response to phage infection, bacteria activate pathways which lead to cell stasis or cell death prior to the release of active virions ^21^. Specifically, a variety of toxin-antitoxin systems have been shown to function as abortive infection systems, where phage infection leads to the formation of active toxin ^19,22–28^. The toxin can then directly interfere with phage development or inhibit essential cellular mechanisms, leading to the eventual death of both phage and host.

Phages in turn have evolved a variety of anti-defense mechanisms which would allow them to thrive in the presence of defense systems ^29–33^. These often include the expression of short ORFs which serve to block the defense system, including the expression of antitoxins that block activated toxins ^25,34–36^.

While defense systems are designed against both lytic and temperate phages, little attention has been given to the way by which these defense systems impact temperate phages and their lysis/lysogeny decisions ^37–39^. Here, we uncover a surprising link between arbitrium-controlled lysis/lysogeny decisions and an anti-phage defense system. We find that the short ORF in the *aimX* gene of the *Bacillus subtilis* phage ɸ3T promotes lysis and codes for a MazE-like antitoxin. Strikingly, we show that MazF activity is crucial for the lysogeny decision, and that ɸ3T becomes almost obligatory lytic in its absence. Additionally, we show that the lytic-induced gene *yosL* encodes for a second MazE-like antitoxin that is conserved in all SPβ-like phages. Finally, in the absence of AimX and YosL, MazEF acts as a non-abortive defense system against a lytic ɸ3T variant, preventing phage propagation. Some functions of MazEF also hold with respect to SPβ-like phages lacking the AimX ORF. Our results suggest that the response of phage ɸ3T to a bacterial defense system is a crucial component of its lysis/lysogeny decision.

## Results

### The lysis/lysogeny decision of phage ɸ3T is affected by the AimX protein

While the activity of the *aimX* gene in SPβ-like phages was previously ascribed to its RNA, it was also noted that phage ɸ3T, where the arbitrium system was initially identified, has a short ORF of 51 amino-acids coded within its *aimX* gene ^3^. We wanted to test whether this ORF plays a role in the lysis/lysogeny decision. To this aim, we cloned this ORF alone (denoted as *aimX*^ORF^, Extended Data Fig. 1) under an IPTG-inducible promoter into an ectopic location in the *B. subtilis* genome. We then monitored infection by a spectinomycin resistance marked ɸ3T strain (used throughout the work unless otherwise noted, to allow measuring lysogenization. methods, Extended Data Fig. 2A) by following the optical density of the population, with and without addition of the AimP arbitrium peptide (SAIRGA, Fig. 1A, Extended Data Fig. 2B). In agreement with previous reports ^3^, we observed a sharp reduction in OD, indicative of lytic infection, followed by growth of lysogens. Lysis was blocked by addition of the AimP peptide to the medium. In contrast, when the AimX ORF was expressed, we found that infection became more lytic (as judged by the extent of reduction in optical density) irrespective of whether AimP was added or not. These results suggest that it is sufficient to express the AimX ORF within *aimX* to promote the lytic fate.

**Figure 1.**
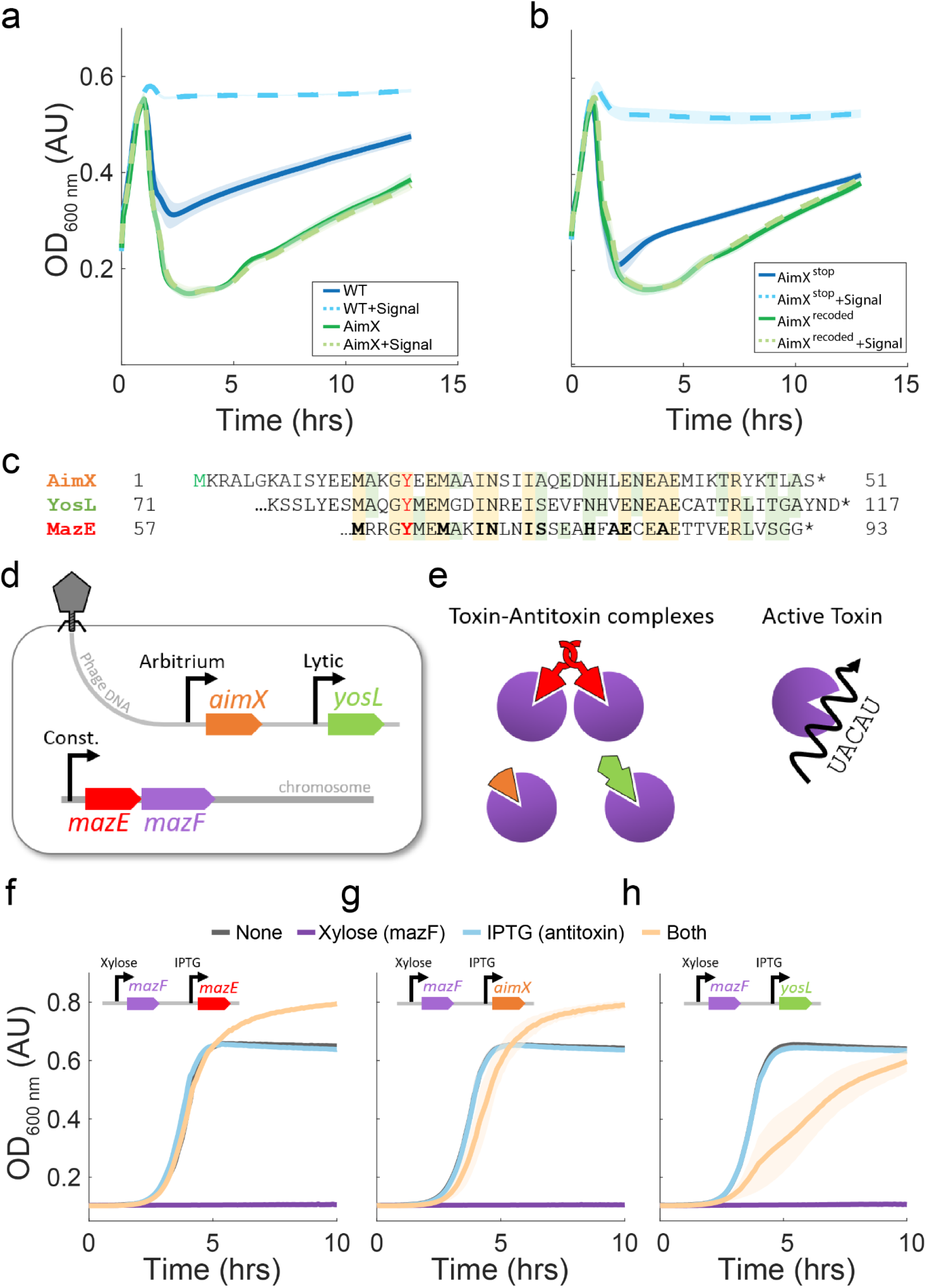
ɸ3T codes for two MazE-like antitoxins, AimX and YosL. (A,B) The *aimX* gene of phage ɸ3T codes for AimX; an antitoxin against the MazF bacterial toxin. Shown are infection growth curves of phage ɸ3T (marked with Spec^R^ gene, methods) infecting different bacterial strains at an initial MOI of 0.1, with (dashed line) or without (solid line) the addition of the mature AimP arbitrium signal. Shaded regions around each curve mark standard error estimation from repeats over three days (methods). (A) Infection of wild-type background strain (PY79;ΔPBSX, blue) and a P^hs^-*aimX*^*ORF*^ strain induced with IPTG (green, note that solid and dashed lines are overlapping). (B) P_hs_-*aimX*^*stop*^ (Blue) and P_hs_-*aimX*^*recoded*^ (Green) both induced with IPTG. IPTG induction was done using a concentration of 100μ M and arbitrium signal (SAIRGA) was added at 10μ M. (C) Protein sequence of ɸ3T AimX and of the homologous parts of ɸ3T YosL and MazE of strain *B. subtilis 168*. Amino-acids shared between the three (yellow) or two (green) are marked. The red-colored Y marks the position mutated to alanine in AimX and YosL. Bold-marked letters in the MazE sequence were deduced to be in contact with the MazF dimer from the crystal structure ^40^. (D) Scheme of the different location and expression patterns of *aimX, yosL* and *mazE* based on published data. (E) Scheme of the proposed interactions between the different antitoxins and MazF and the impact of MazF in absence of antitoxins. (F-H) MazE, AimX and YosL counteract MazF toxicity. Results for three strains all harboring a Δ*mazEF* deletion and coding for a xylose inducible *mazF* (P_xyl_-*mazF*). Each of the strains codes for an IPTG-inducible putative antitoxin (F) P_hs_-*mazE*; (G) P_hs_-*aimX*; (H) P_hs_-*yosL*. The four conditions used are – no inducers (green), IPTG (100μ M in E,F and 30μ M in G, blue), xylose (2% w/v; red) and both inducers (cyan).

To further demonstrate that the effect of the AimX ORF is exerted by the translated protein and not by its RNA transcript, we constructed two additional AimX-coding variants under IPTG control (Figs. 1B, 1C, Extended Data Fig. 2B). In the first strain, AimX^stop^, we mutated the first methionine codon of AimX to a stop codon. In the second, AimX^recoded^, we re-coded the AimX transcript using different codons to obtain the same translated protein, with a different RNA sequence (Extended Data Fig. 1). Infecting these strains with ɸ3T, either with or without the addition of AimP, we found that the AimX^stop^ mutant had no effect on infection, while AimX^recoded^ had the same effect as the AimX^ORF^ strain. These results strongly suggest that the function of the ɸ3T *aimX* gene is mediated by the AimX protein.

### ɸ3T proteins AimX and YosL are both antitoxins of the bacterial MazF toxin

Following the identification of AimX as a protein important for the lysis/lysogeny decision, we wondered what its function is. AimX has been previously annotated as a homolog of the C-terminal part of YosL, which is coded by many *Bacillus* phages and is common to all SPβ-like phages (Fig. 1C) ^3,41^. In both ɸ3T and SPβ, *yosL* is encoded within an operon containing phage replication-related genes, suggesting that it is expressed during the lytic cycle (Fig. 1D). Indeed, *yosL* of the SPβ prophage is expressed only during prophage induction ^41–43^. Searching for further homology using BLAST, we found that both AimX and YosL are weakly homologous to MazE (Fig. 1C), the antitoxin protein of the bacterial host-encoded MazEF toxin-antitoxin system. Specifically, AimX and the C-terminal part of YosL are homologous to the C-terminal part of MazE, which is the MazF binding site, sharing a large number of the residues on the MazF binding surface (56% and 52% sequence identity for YosL and AimX, respectively) ^40^. AimX and YosL lack the N-terminal dimerization domain of MazE, but YosL has an alternative unrelated domain in its N-terminus (Fig. 1E) ^40,44^.

If AimX and YosL are functionally homologous to MazE, we would expect them to deactivate the toxicity of MazF RNA endoribonuclease activity (Fig. 1E) ^40,44,45^. To experimentally study this hypothesis, we constructed three strains. Each of the strains is deleted for the *mazEF* operon and codes for *mazF* under a xylose inducible promoter. The strains additionally code for either *aimX*^*recoded*^, *yosL* or *mazE* under an IPTG-inducible promoter (Fig. 1F-H). When MazF was expressed on its own by addition of xylose during early growth, it was able to halt growth completely, while co-expression of MazE suppressed MazF toxicity, as previously described (Fig. 1F, the extended growth is explained by the addition of xylose, Extended Data Fig. 2C) ^40,44^. In accordance with its proposed function as an antitoxin, we found that expression of AimX together with MazF also allowed for normal growth, while expression of AimX on its own had no effect on growth (Fig. 1G). These results fit the definition of interactions between a toxin and its antitoxin.

To further validate the similarity of MazE and AimX, we mutated the tyrosine at residue 18 of AimX to alanine. This residue is conserved between the two proteins and was shown to be crucial for MazE-MazF interactions (Fig. 1C) ^40^. In accordance with its putative role in AimX function, we found that expression of AimX^Y18A^ had no effect on the toxicity of MazF (Extended Data Fig. 3A).

Repeating the experiments with YosL, we found that modest expression of *yosL* also partially blocked MazF toxicity, as expected from a MazE homologue (Fig. 1H). However, in contrast to both AimX and MazE, *yosL* overexpression was toxic to the cells in the absence of MazF, suggesting that YosL has an additional function independent of blocking MazF activity and explaining the partial growth of a strain expressing YosL and MazF (Extended Data Fig. 3D). Mutation of the conserved tyrosine of *yosL* (position 82, Fig. 1C) to alanine eliminated YosL ability to block MazF toxicity (Extended Data Fig. 3B), but did not eliminate its independent toxicity at high expression levels (Extended Data Fig. 3D). YosL therefore serves as a MazE-like antitoxin but has an additional, independent, function. As YosL has an additional domain, we wondered whether expression of the MazE-homologous C-terminal domain of yosL would not show the toxicity effect. Indeed, we found that expression of the C-terminal domain blocked MazF toxicity without inducing any lethal effects on its own (Extended Data Fig. 3C,F).

Finally, we looked at the predicted model structures of the complexes of AimX and YosL with MazF dimers (Extended Data Fig. 4). To do this, we first predicted the structures of the MazE-homologous parts of AimX and YosL by performing homology modeling. Then, we superimposed each of the predicted structures onto MazE in the solved crystal structure of MazE and MazF ^40^. These models suggest that despite the sequence differences between AimX/YosL and MazE, the three antitoxins still form similar complexes with the MazF dimer, in which some of the binding interactions are equivalent. In summary, AimX and YosL of phage ɸ3T are homologous both in sequence, predicted structure and function to the MazE antitoxin.

### MazF activity is required for lysogenization in ɸ3T

The above results suggest a link between the lysis/lysogeny decision of phage ɸ3T and the MazEF system. To study this, we tested the effect of Δ*mazF* and Δ*mazEF* deletion mutants on phage ɸ3T lysis/lysogeny decisions (Fig. 2, Extended Data Fig. 2C).

**Figure 2.**
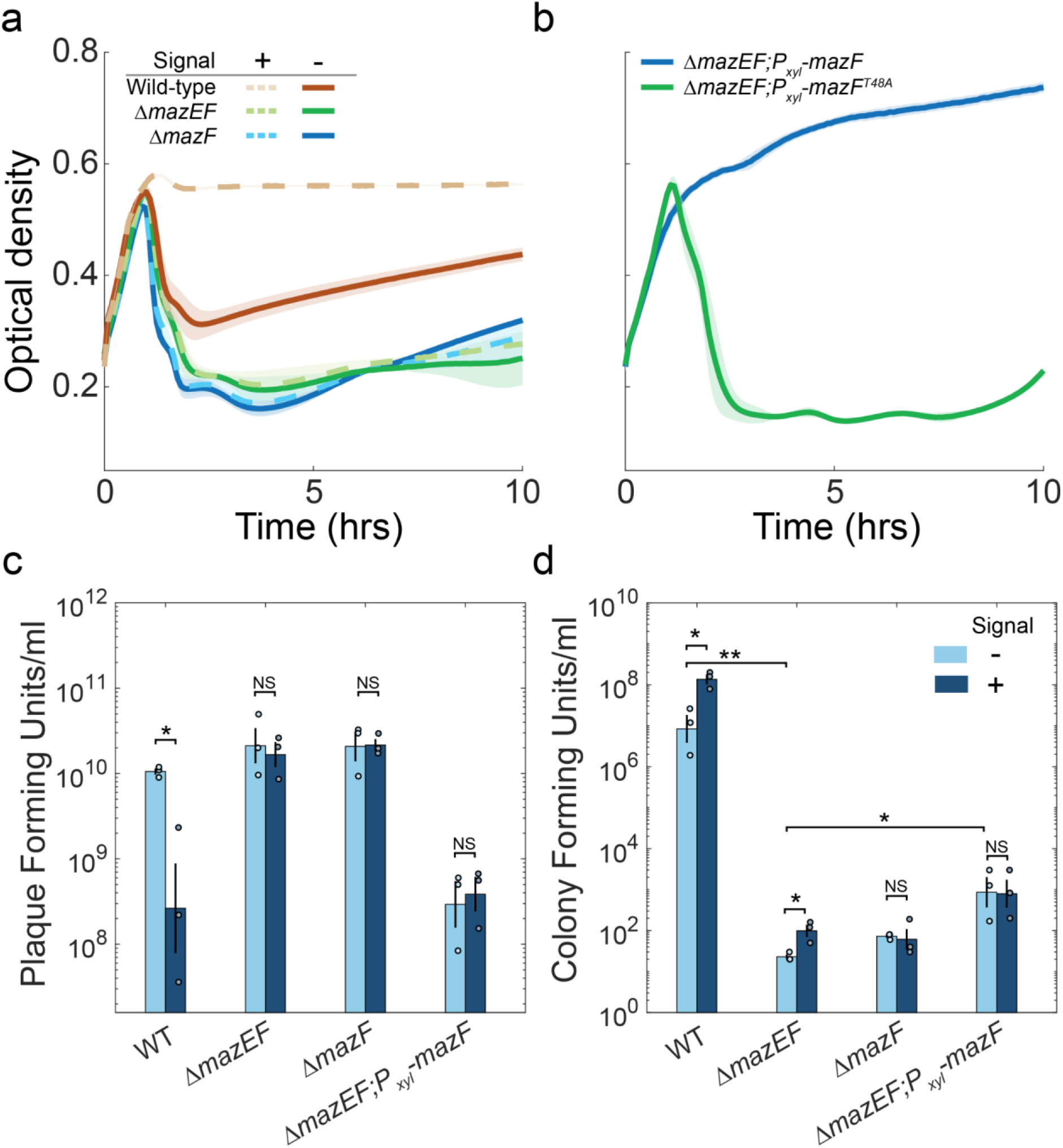
MazF is necessary for ɸ3T lysogenization. (A) growth curves of different strains infected by ɸ3T at MOI of 0.1, either in the absence (solid line, darker color) or presence (dashed line, lighter color) of the arbitrium peptide (SAIRGA). The following strains were used; wild-type (red), Δ*mazF* (blue), Δ*mazEF* (green). (B) growth curves of different strains infected by ɸ3T at MOI of 0.1. The following strains were used; Δ*mazEF;P*_*xyl*_*-mazF* (green), Δ*mazEF;P*_*xyl*_*-mazF*^*T48A*^ (blue, catalytic dead mutation). Both strains were induced with 2% xylose. (C,D) Free phage and lysogen numbers were measured for infection of various bacterial strains (see legends) by a spectinomycin marked ɸ3T phage. (C) Plaque forming units (PFU) were measured after an overnight infection by plating on a Δ*mazEF* indicator strain. (D) lysogens were enumerated by measuring colony forming units on spectinomycin plates. All measures were done in triplicates. Shaded regions around growth curves in A,B mark standard error between biological repeats. Error bars in C,D mark logarithmic standard error.

We first followed infection by its impact on cellular growth (Fig. 2A). We found that ɸ3T infection of the two mutants was more lytic than its infection of the wild-type. Infection of the mutants was also insensitive to the addition of the ɸ3T AimP peptide (Fig. 2A). The above results were also reflected in plaque counts after an overnight infection (Fig. 2C), with a 80-fold reduction in total PFU upon addition of the arbitrium signal to the wild-type (n=3, p=0.03, two-sided t-test), but not upon its addition to the Δ*mazF* and Δ*mazEF* mutants (n=3, p=0.93,0.7 respectively, two-sided t-test). To further validate the importance of MazF to the lysogeny pathway, we used a Δ*mazEF* deletion strain expressing *mazF* under a xylose-inducible promoter and assayed infection with xylose added to the medium at the time of infection. We found that addition of xylose strongly represses lytic infection as reflected by both the growth curve (Fig. 2B) and plaque counts (Fig. 2C, n=3, p=0.005, two-sided t-test). Notably, the induction of MazF at the mid-log stage (OD_600_ of 0.3) did not lead to immediate cessation of growth in the absence of phage infection (Extended Data Fig. 5A), though measured CFU levels were reduced 10-fold (Extended Data Fig. 5B).

MazF RNA ribonuclease activity is known to depend on the threonine residue at position 48^40^. We found that xylose-dependent expression of MazF^T48A^ did not show any effect on infection dynamics in a Δ*mazEF* background, confirming the importance of MazF ribonuclease activity for its function in suppression of lysis (Fig. 2B).

To directly measure the importance of MazEF for lysogenization, we infected the wild-type, Δ*mazF* and Δ*mazEF* strains with the marked ɸ3T either with or without the arbitrium peptide and plated infected cells 20 minutes and two hours after infection on spectinomycin plates, to select for lysogens (Fig. 2D, Extended Data Fig. 6). We found that the number of wild-type lysogens increased more than 10-fold when arbitrium was added to the medium, as expected (n=3, p=0.03, two-tailed t-test). In contrast, the number of lysogens on Δ*mazF* and Δ*mazEF* background was reduced by more than five orders of magnitude compared to that of the wild-type (n=3, p<2✕10^−4^ for both, two-tailed t-test). The number of lysogens on Δ*mazF* did not show any dependence on the arbitrium signal (n=3, p=0.8, two-tailed t-test), while the number of lysogens on a Δ*mazEF* background showed a weak (4-fold) but statistically significant dependence on addition of the arbitrium signal (n=3, p=0.02, two-tailed t-test). We also measured the formations of lysogens upon infection of a Δ*mazEF;P*_*xyl*_*-mazF* strain with xylose added (methods). We found that MazF expression led to a 40-fold increase in lysogen numbers compared to the Δ*mazEF* background (n=3, p=0.01, two-tailed t-test). Altogether, our results suggest that the core MazEF system of *B. subtilis*, and the ribonuclease activity of MazF specifically, is necessary for phage ɸ3T lysogenization and for the function of the arbitrium system.

### MazF activity inhibits prophage induction

It was previously shown that the arbitrium system also regulates prophage induction in SPβ-like phages. Specifically, prophage induction is promoted by DNA damage and suppressed by the arbitrium peptide ^4–6^. We therefore wondered whether MazEF affects prophage induction. To address this question we isolated one of the rare lysogens formed upon infection of Δ*mazEF* by ɸ3T. We found that these lysogens were stable during growth, suggesting that MazEF is not necessary for maintenance of the lysogenic state. The Δ*mazEF* lysogen was able to induce upon addition of mitomycin C (MMC), a potent DNA damaging agent, and yield phenotypically wild-type phages in terms of their infection pattern in wild-type and ΔmazEF mutant. Monitoring cell density during prophage induction (Extended Data Fig. 7A), we found that these lysogens were able to induce in a similar fashion to wild-type lysogens. However, in contrast to the wild-type, Δ*mazEF* lysogen prophage induction was insensitive to the addition of the AimP peptide. In addition, we isolated lysogens in a Δ*mazEF*;P_xyl_-*mazF* background and induced them with MMC either with or without the expression of *mazF* through addition of xylose. We found that expression of *mazF* prevented prophage induction and cell lysis (Extended Data Fig. 7B). These results suggest that similar to its function during infection, the arbitrium system also controls prophage induction by modulating the MazEF system.

### MazEF partially modifies phage SPβ lysis/lysogeny decision

Many SPβ-like phages, and specifically phage SPβ itself, do not code for any clear open reading frame within their *aimX* locus. However, all of these phages code for *yosL*^3,41^. It is therefore unclear whether the MazEF system still impacts the lysis/lysogeny decision of these phages. To explore this question, we first examined the impact of a *ΔmazEF* mutation on SPβ infection dynamics. Similarly to ɸ3T, we find that infection in a *ΔmazEF* tends more towards lysis than that of the wild-type, and becomes insensitive to addition of the SPβ arbitrium signal (GMPRGA, Extended Data Fig. 8A,B). Notably, and in contrast to ɸ3T, this was not the case during prophage induction where the mutant still responded to addition of the signal (Extended Data Fig. 8C). Additionally, in contrast to ɸ3T, we did not observe a marked reduction in lysogen numbers during infection (assayed using a spectinomycin resistance marked phage ^46^) in the *ΔmazEF* compared to the wild-type (Extended Data Fig. 8D, n=3, p=0.34, two-tailed t-test). These results suggest that MazEF has some effects on SPβ lysis, but that lysogenization is not dependent on MazF activity.

### In the absence of phage anti-toxins, MazEF serves as a defense system against lytic ɸ3T

Several toxin-antitoxin systems are known to act as anti-phage defense systems ^19,22–28^. MazEF of *E. coli* has also been attributed as such a defense system (^47,48^, but c.f, ^49^) and a MazF homolog containing protein serve to defend against *S. aureus* phages ^20^. The expression of phage-encoded antitoxins is a known defense-evasion strategy of phages ^25,34^. We therefore wondered whether MazEF acts as a defense system that phage ɸ3T overcomes by expressing *aimX, yosL* or both. To address this hypothesis, we constructed an obligatory lytic unmarked mutant, ɸ3T(vir), by deleting the *yopR* repressor ^4,41^. This strain could be maintained and modified in a lysogenic state in a background strain coding for an IPTG-inducible *yopR* (methods). We further introduced into the ɸ3T(vir) background an *aimX* deletion, *yosL* deletion or both (methods). We then assayed the four resulting strains for their ability to infect the wild-type or Δ*mazEF* mutant. To this end, we monitored plaque forming units after overnight infections of each of the phages on either strain at multiplicity of infection (MOI) of 0.1 (Fig. 3A). We find that while *aimX* deletion did not lead to considerable reduction in PFU levels on either strains, deletion of *yosL* from ɸ3T(vir) led to a small (5-fold) but statistically significant reduction when infecting the wild-type compared to infection of Δ*mazEF* (n=3, p=0.02, two-tailed t-test). Deletion of both *aimX* and *yosL* showed strong synergism, leading to a reduction of more than 6 orders of magnitude in PFU on the wild-type compared to the Δ*mazEF* (n=3, p<10^−3^, two-tailed t-test). This was further corroborated by monitoring infection dynamics of the four strains on wild-type and Δ*mazEF* strains (Extended Data Fig. 9A-C). These results suggest that in the absence of *aimX* and *yosL* phage ɸ3T cannot proceed with the lytic process. This may arise from a defense function of the MazEF system, or alternatively, from a miss-regulation of the lysis/lysogeny decision due to the phage *aimX* deletion. However, the necessity of *yosL* deletion, which is expressed during phage replication, does not support the latter hypothesis.

**Figure 3.**
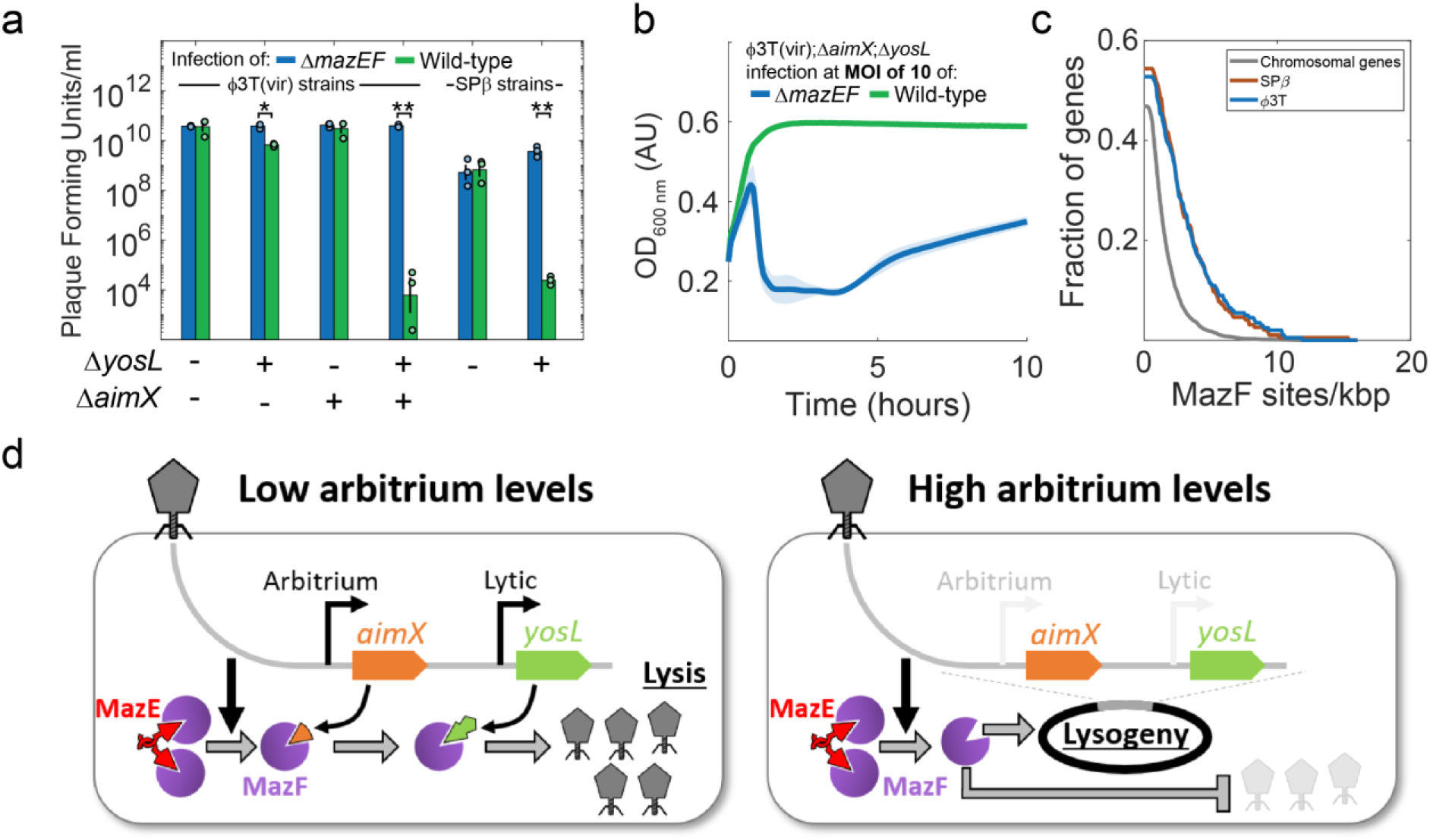
Deletion of phage MazE-like antitoxins renders a virulent ɸ3T strain and phage SPβ sensitive to MazEF activity without eliciting cell death. (A) PFU after overnight infection at MOI of 0.1 of four ɸ3T(vir) strains and two SPβ strains on either wild-type (green) or Δ*mazEF* (blue) bacterial strains. Strain genotypes are described on the x-axis. - indicates that the gene is not deleted and + indicates that the gene is deleted. Only *yosL* status is marked for phage SPβ, as *aimX* is irrelevant. (B) growth curves during infection by phage ɸ3T(vir)*;aimX;yosL* at MOI of 10 on wild-type (green) and Δ*mazEF* (blue) strains. (C) Fraction of genes with a density of putative MazF cleavage per kilobasepair higher than the density denoted on the x-axis. Shown is the data for *B. subtilis* 168 chromosome (gray, excluding SPβ), SPβ (brown) and ɸ3T (blue). Note that the fraction of genes with 0 DNA cleavage sites is equal to one minus the value of the function at x=0. (D) A schematic model of the lysis/lysogeny process. In the absence of arbitrium signal (left), AimX and later YosL inhibits MazF and allows the lytic cycle to commence. In the presence of an arbitrium signal, MazF remains active, induces lysogeny and prevents lysis if the antitoxins are not expressed.

To further strengthen the assertion that MazEF acts as a defense system, we turned to phage SPβ which, as mentioned, encodes only for the YosL antitoxin and where MazEF does not affect lysogenization. We deleted *yosL* from a spectinomycin marked SPβ, and found that this strain became sensitive to MazEF defense without significantly affecting lysogenization (Fig. 3A, Extended Data Fig. 10A,B). These results further suggest that MazEF can act as defense system against SPβ-like phages in absence of phage-expressed antitoxins, irrespective of its role in lysis/lysogeny decisions.

Toxin-antitoxin systems are thought to act as abortive infection systems, though recently it was shown that they may act preferentially on phage components ^25,49^. To check whether MazEF acts as an abortive infection system, we infected the wild-type and Δ*mazEF* strains with the ɸ3T(vir);*aimX*;*yosL* mutant at a MOI of 10 to ensure infection of the majority of cells. We then followed the infection on both wild-type and Δ*mazEF* strains. We found that infection of Δ*mazEF* led to rapid reduction in cell density, while infection of the wild-type did not lead to any cell death and growth was normally maintained (Fig. 3B). These results were corroborated by plaque and colony counts two hours after infection (Extended Data Fig. 9D,E). Altogether, these results suggest that the MazEF system defends against the mutant phage infection but does not act as an abortive infection system, as it has no discernible impact on cell viability.

### Transcripts of Phages ɸ3T and SPβ are enriched in MazF cleavage sites

One potential explanation for the ability of MazF to efficiently eliminate the antitoxin deficient virulent variants of ɸ3T and SPβ without causing bacterial dormancy is that their RNA is more sensitive to MazF activity than that of the bacterium. The *B. subtilis* MazF cleavage site has been characterized to be ‘UACAU’, though some deviations from this consensus are tolerable ^45^. To understand the relative sensitivity of ɸ3T and *B. subtilis* derived RNAs to MazF, we calculated the number of cleavage sites per kb in each of the genes of strain 168 (excluding the SPβ phage) and for phages ɸ3T and SPβ. To account for the %GC difference between the phage genomes and the bacterial chromosome, we also calculated the normalized deviation of the real number of cleavage sites per gene from the random number of sites in a reshuffled gene (maintaining the same %GC content, methods). We find that both phages had a significantly higher proportion of genes with a high density of cleavage sites using both measures. (Fig. 3C, Extended Data Fig. 11). For example, 17.5% of genes in SPβ and ɸ3T had a cleavage site density of 4 sites per kb or more, compared to just 3.5% of chromosomal genes (p<10^−45^ based on resampling of the chromosomal distribution, methods).

## Discussion

Here, we identified an intimate interaction between the arbitrium communication system of phage ɸ3T and the MazEF toxin-antitoxin system. AimX, the arbitrium lysis actuator, is a MazE homolog. AimX serves to block MazF anti-phage activity (together with the later expressed gene YosL) allowing lysis, while uninterfered MazF activity triggers lysogeny by a mechanism that remains to be elucidated. Finally, we showed that the MazEF system acts as a defense system against SPβ and a lytic ɸ3T mutant in the absence of their respective antitoxin genes.

Defense mechanisms work against temperate phages, but many of them are activated only during the lytic cycle and therefore are not expected to interfere with lysogeny ^18,21^. Two possible counter-defense strategies of temperate phages are therefore to initiate lysogeny or to express anti-defense genes. Here we found that the arbitrium system of phage ɸ3T controls the execution of either of these strategies, depending on concentration of the arbitrium signal. To this aim, ɸ3T co-opted MazF activity as a signal for initiation of lysogeny. This may occur by specific cleavage events which remodel or reduce the levels of a specific pro-lytic transcript, or by a more general sensing mechanism which initiates lysogeny upon blocking of the lytic pathway. Notably, this function is specific for ɸ3T and is not found in SPβ, stressing the importance of local adaptations within this clade, which probably co-evolved with the acquisition of the AimX antitoxin gene. More generally, lysogeny may serve as a counter-defense response in other phages and it would be intriguing to identify additional defense systems involved in the lysis/lysogeny decision.

In many arbitrium-coding phages out of the SPβ family, the *aimX* locus is an antisense to a short XRE-domain transcription factor which can serve as the phage repressor, homologous to the λ phage cI repressor. RNAseq experiments suggest that aimX functions as a non-coding RNA that may reduce repressor transcript levels upon expression and promote lysis^13^. Notably, some of these *aimX* transcripts also code for small ORFs, but their function has not been determined experimentally.

The MazEF system which we identified here to restrict infection of mutant SPβ and ɸ3T phages is atypical for a toxin-antitoxin defense system. First, we find that in contrast to previously identified phage defense toxin-antitoxin systems, this system is non-abortive^23,25,50^. The ability of this system to specifically restrict phage activity without affecting cellular growth may partially depend on the excess of UACAU cleavage sites found in the (generally low GC) phage transcripts of *B. subtilis* compared to the chromosomal genome (Fig. 3C, Extended Data Fig. 11). As reported in previous studies, MazEF activation is also not unique to phage infection and it is activated under other physiological conditions, though at least some of the conditions may relate to prophage induction ^51^. Notably, the ToxIN phage defense system also does not directly kill the cell and it would be interesting to study its possible effect on temperate phages ^49^. Second, in contrast with the typical accessory nature of phage defense islands ^15,16,19,20,52^, the MazEF system is part of the core genome of *B. subtilis* ^53^. Third, we show here that MazEF can both serve as a defense system but also has a regulatory role. While we cannot rule out completely that MazF activity prevented phage propagation simply by leading to a regulatory failure in the mutant phages, we find this to be less likely - in phage SPβ YosL has no regulatory role in lysis/lysogeny and yet is critical for productive infection in the presence of MazEF. It would be interesting to understand whether other seemingly non-phage defense related toxin-antitoxin systems play a role in bacteria-phage interactions.

We propose a model for the phage-host interaction, which includes the following processes (Fig. 3D): i) Phage infection activates the MazEF system. iia) Under low AimP signal levels, ɸ3T counteracts MazF ribonuclease activity by expressing AimX and initiating the lytic pathway. IIIa) During lysis, YosL is also expressed to further inhibit MazF defense. iib) In contrast, under high AimP signal levels, AimX is not expressed and the phage switches into a lysogenic state through a MazF-dependent process. iiib) Following lysogenization, MazF becomes inactive. Many parts of this model are still missing, including the activation and deactivation signals for MazF, the activation of lysogeny by MazF and the exact nature of its phage defense activity.

Finally, lysogeny in both SPβ and ɸ3T is maintained and regulated by the *yop* operon which is located immediately downstream of *aimX* ^4^. While AimX is found in the ɸ3T-like subset of the SPβ-like family, other lysogeny related genes and specifically the *yopM-R* operon is conserved throughout the entire family and is expected to be regulated by the arbitrium system. We therefore anticipate that an additional arbitrium-dependent lysis/lysogeny mechanism works in SPβ and other phages not coding for AimX, and may also work in parallel to the MazEF system in ɸ3T and related phages. Further work is needed to untangle the intricate cross-talk between the *yopM-R* operon, the arbitrium system and the MazEF system.

## Methods

### Strain construction

All bacterial strains, plasmids and primers used in this study are listed in Supplementary Tables 1-3. To construct new *B. subtilis* strains, standard transformation protocols were used for genomic integration and plasmid transformation ^54^. The *B. subtilis* lab strain PY79 contains an additional DNA damage-induced lytic element (the PBSX prophage-derived bacteriocin) and therefore we used as a common background a markerless *Δxpf* strain in which PBSX cannot induce ^55^.

*mazF, yopR*(SPβ) *yosL(SPβ)* were deleted by transformation of a kanamycin resistance cassette from a *Bacillus subtilis* 168 deletion library ^56^ into the appropriate strains. The kanamycin resistance cassette was then excised using a Cre/*lox* system ^56^. For the *mazEF, yosL(ɸ3T), yopR(ɸ3T) and aimX(ɸ3T)* deletions, we used a long flanking homology PCR method to replace genes with either a kanamycin or an erythromycin resistance cassette ^6^. To delete phage repressors *yopR*(SPβ) and *yopR(*ɸ3T) we first introduced a Phs-*yopR*(SPβ) \ Phs-*yopR*(*ɸ3T*) construct into a lysogen strain. All primers used are listed in Supplementary Table X.

Marked ɸ3T was constructed by insertion of spectinomycin resistance(spec) between the convergently transcribed genes *yolB* and *yolC* using a long flanking homology PCR method. Similarly to SPβ:spec ^46^, spectinomycin resistance was inserted in such a way that a terminator is located on each side of it.

### Plasmid construction

pAEC2220 was constructed by amplifying the *aimX* gene from genomic DNA of a ɸ3T lysogen (AES7006) using the POL168 and POL170 primer pair. The PCR product was digested with SacI-HF and NheI-HF, and cloned into ECE174 digested with NheI-HF and SacI-HF.

pAEC2571 was constructed by amplifying the whole pAEC2220 using the POL314 and POL315 primer pair that introduced a point mutation of start codon to stop codon. The original plasmid in the PCR product was degraded using DpnI. The PCR product was closed after adding phosphate using PNK enzyme, followed by ligation.

pAEC2229 was constructed by cloning a pre-ordered gBlocks™ Gene Fragments (IDT, Israel) of recoded aimX (Extended Data Fig. 1) into ECE174, both digested with NheI-HF and SacI-HF.

pAEC2513 was constructed by amplifying yopR(ɸ3T) gene from genomic DNA of a ɸ3T lysogen (AES7006) using the POL279 and POL280 primer pair. The PCR product was digested with SacI-HF and NheI-HF, and cloned into ECE174 digested with NheI-HF and SacI-HF.

pAEC2610 was constructed by amplifying the whole pAEC2229 using the SOB670 and SOB671 primer pair that introduced a point mutation Y18A. The original plasmid in the PCR product was degraded using DpnI. The PCR product was closed after adding phosphate using PNK enzyme, followed by ligation.

pAEC2550 was constructed by amplifying pDR111 using the SOB627 and SOB628 primer pair and amplifying mazE gene from *B*.*subtilis* genomic DNA using the SOB629 and SOB630 primer pair. The purified DNA fragments were then ligated using the Gibson assembly protocol ^57^.

pAEC2246 was constructed by amplifying pDR111 using the SOB466 and SOB467 primer pair and amplifying yopR gene from an SPβ lysogen genomic DNA (AES7975) using the SOB476 and SOB477 primer pair. The purified DNA fragments were then ligated using the Gibson assembly protocol ^57^.

pAEC2539 was constructed by amplifying yosL gene from genomic DNA of a ɸ3T lysogen (AES7006) using the POL295 and POL296 primer pair. The PCR product was digested with SacI-HF and NheI-HF, and cloned into ECE174 digested with NheI-HF and SacI-HF.

pAEC2573 was constructed by amplifying the whole pAEC2539 using the POL318 and POL319 primer pair that introduced a point mutation Y82A. The original plasmid in the PCR product was degraded using DpnI. The PCR product was closed after adding phosphate using PNK enzyme, followed by ligation.

pAEC2276 was constructed by amplifying the mazF gene from genomic DNA of *B*.*subtilis* 168 using the POL256 and POL257 primer pair. The PCR product was digested with SpeI-HF and BamHI-HF, and cloned into ECE137 digested with SpeI-HF and BamHI-HF.

pAEC2604 was constructed by amplifying the whole pAEC2276 using the POL312 and POL313 primer pair that introduced a point mutation T48A. The original plasmid in the PCR product was degraded using DpnI. The PCR product was closed after adding phosphate using PNK enzyme, followed by ligation.

pAEC2615 was constructed by amplifying pAEC2539 using the SOB676 and SOB677 primer pair, to delete yosL residues 2-70. The original plasmid in the PCR product was degraded using DpnI. The PCR product was closed after adding phosphate using PNK enzyme, followed by ligation.

### Growth media and conditions

All the experiments in this study were performed in rich media-Lysogeny Broth (LB): 1% tryptone (Difco), 0.5% yeast extract (Difco), 0.5% NaCl. Liquid cultures were grown with shaking at 220 RPM and a temperature of 37°. When preparing plates, medium was solidified by addition of 2% agar. Antibiotics were added (when necessary) at the following concentrations: spectinomycin: 100 µg ml^-1^, chloramphenicol: 5 µg ml^-1^, kanamycin: 10 µg ml^-1^, MLS: 3 µg ml^-1^ erythromycin + 25 µg ml^-1^ lincomycin. For experiments that involve infection, media were supplemented with 0.1 mM MnCl_2_ and 5 mM MgCl_2_. The Phs promoter was activated with 100 μM isopropyl-ß-D-thiogalactopyranoside (IPTG) unless stated otherwise, and the Pxyl promoter was activated with 2% (w/v) xylose.

### Plaque forming assay

Samples for PFU measurements were collected from cultures centrifuged for 5 min at 4,000 r.p.m at room temperature. Next, the supernatant was filtered using a 0.2 μm filter (Sartorius Stedim biotech cat. 14-555-270). 100 μL of filtered supernatant at an appropriate dilution was then mixed with 200 μL of *B. subtilis* wild-type or *ΔmazEF* indicator strain grown to OD_600_ in MMB (LB supplemented with 0.1 mM MnCl_2_ and 5 mM MgCl_2_), and left to incubate at room temperature for 5 minutes. Three ml of molten LB-0.5% agar medium (at 60°C) supplemented with 0.1 mM MnCl_2_ and 5 mM MgCl_2_ was added, mixed, and then quickly overlaid on LB-agar plates. Plates were then incubated at 37° for 1 hour and then ON at room temperature to allow plaques to form.

### Growth dynamics

To examine growth dynamics in different conditions, strains were grown overnight in LB at 37° with shaking at 220 RPM then diluted by a factor of 1:100 into fresh LB media and grown upon reaching OD_600_ = 0.3. For prophage induction experiments, cultures were then supplemented with 0.5 µg ml^-1^ of MMC, 10 µM of peptides and 2%(w\v) xylose when indicated. For infection experiments, cultures were then infected with SPβ or ɸ3T and supplemented with 10 µM of peptides, 2%(w\v) xylose and 100µM IPTG when indicated. For toxicity experiments, cultures were then subsequently diluted by a factor of 1:1000 into fresh LB media supplemented with 2%(w\v) xylose and IPTG, as indicated. Optical density measurements at a wavelength of 600 nm were performed in a 96-well plate using a plate reader (SPARK® multimode microplate reader, Tecan, USA).

### Infection experiments

Strains were grown overnight in LB at 37° with shaking at 220 RPM then diluted by a factor of 1:100 into fresh LB media. Upon reaching OD600 = 0.1 cultures were supplemented with 10 µM of peptides and 2%(w\v) xylose as indicated. Strains were then grown at 37° with shaking at 220 RPM to OD_600_ = 0.3 and infected with SPβ or ɸ3T phages harboring a spectinomycin resistance marker at an MOI of 0.1. For quantification of lysogens, 1ml of infected cultures were taken 20 minutes and 2 hours after infection. Cells were washed 3x with 1 ml PBS to remove unbound phage and then diluted and spread on LB plates with spectinomycin to select for cells that had become lysogenized.

For PFU quantification, 1 ml of infected cultures was collected ON, centrifuged for 5 min at 4,000 r.p.m at room temperature and filtered using a 0.2 μm filter. Then, a plaque forming assay was performed.

### Cleavage site statistics

Sequence and annotation of *B. subtilis* str. 168 (accession NC_000964.3) and ɸ3T(accession KY030782) are based on their genbank annotation files. For each of the genes in the annotations we calculated the number of MazF consensus cleavage sites (UACAU) in the gene’s ORF and divided by gene length to obtain the cleavage sites density. For each gene we also ran a 1000 random shuffling of the ORF sequence and calculated the mean and standard deviation of the number of cleavage sites in this random set. We calculated the normalized deviation of the gene number of cleavage sites as the number of sites minus the average on the shuffled set and divided this by the standard deviation. As there are ∼200 genes in SPβ and ɸ3T and ∼4000 in strain 168 (excluding SPβ) we calculated the probability of obtaining a certain cumulative value in the phages vs. the chromosome by running 1000 random sampling of 200 genes from the chromosome and calculating the value for each of these runs. The p-value for obtaining the value for the phage was then calculated based on this subsample.

### Homology modeling

Modeller ^58^ was used to predict the structures of YosL (positions 78-104) and AimX (positions 14-40) by homology modelling, based on their sequence similarity to MazE (which was validated with ConSurf using default settings and 300 hundred sequences^59,60^). The MazE-YosL and MazE-AimX sequence identities were 56% and 52%, respectively. Each of the modelled structures was then superimposed separately onto MazE in its complex structure with MazF (PDB ID: 4me7). Such superimposition often involves steric clashes with surrounding residues. Therefore, the entire YosL/AimX-MazF_2_ complex was computationally refined using the GalaxyRefineComplex web server ^61^, to relieve at least some of these clashes.

## Acknowledgements

We thank Shaul Pollak, Joshua Jones, Adi Stern, David Burstein and members of the Eldar lab for fruitful discussions and comments on the manuscript and Nir Ben-Tal for suggestions about structural modeling. The Eldar lab is funded by a European Research Council grant 724805 and by Israel Science Foundation grant 2228/21. The funders had no role in study design, data collection and analysis, decision to publish or preparation of the manuscript.

## Author Contributions Statement

PG, SOB, NA and AE were involved in the conceptualization of the study. PG and SOB performed the experiments. PG, SOB, NA, AK and AE analyzed the data and formulated the theoretical predictions. PG, SOB, NA and AE wrote the original draft. PG, SOB, NA, AK and AE made the figures. PG, SOB, NA and AE edited the manuscript.

## Supplementary figures and files

This manuscript contained 11 Extended data figures, all attached at the end of the manuscript. One zipped supplementary file contain 16 excel files with all experimental data used in this work and a short Word file describing the content of the excel files.

## References

1. Brady, A. et al. Molecular Basis of Lysis–Lysogeny Decisions in Gram-Positive Phages. Annu. Rev. Microbiol. (2021) doi:10.1146/annurev-micro-033121-020757.

2. Oppenheim, A. B., Kobiler, O., Stavans, J., Court, D. L. & Adhya, S. Switches in bacteriophage lambda development. Annual review of genetics 39, 409–429 (2005).

3. Erez, Z. et al. Communication between viruses guides lysis-lysogeny decisions. Nature 541, 488–493 (2017).

4. Brady, A. et al. The arbitrium system controls prophage induction. Current Biology 31, 5037–5045. e3 (2021).

5. Bruce, J. B., Lion, S., Buckling, A., Westra, E. R. & Gandon, S. Regulation of prophage induction and lysogenization by phage communication systems. Current Biology 31, 5046–5051. e7 (2021).

6. Aframian, N. et al. Dormant phages communicate via arbitrium to control exit from lysogeny. Nature microbiology 7, 145–153 (2022).

7. Aframian, N. & Eldar, A. A Bacterial Tower of Babel: Quorum-Sensing Signaling Diversity and Its Evolution. Annual Review of Microbiology 74, 587–606 (2020).

8. Neiditch, M. B., Capodagli, G. C., Prehna, G. & Federle, M. J. Genetic and Structural Analyses of RRNPP Intercellular Peptide Signaling of Gram-Positive Bacteria. Annual Review of Genetics 51, 311–333 (2017).

9. Wang, Q. et al. Structural basis of the arbitrium peptide–AimR communication system in the phage lysis–lysogeny decision. Nature microbiology 3, 1266–1273 (2018).

10. Del Sol, F. G., Penades, J. R. & Marina, A. Deciphering the molecular mechanism underpinning phage arbitrium communication systems. Molecular cell 74, 59–72. e3 (2019).

11. Guan, Z. et al. Structural insights into DNA recognition by AimR of the arbitrium communication system in the SPbeta phage. Cell Discovery 5, 29 (2019).

12. Pei, K., Zhang, J., Zou, T. & Liu, Z. AimR Adopts Preexisting Dimer Conformations for Specific Target Recognition in Lysis-Lysogeny Decisions of Bacillus Phage phi3T. Biomolecules 11, 1321 (2021).

13. Stokar-Avihail, A., Tal, N., Erez, Z., Lopatina, A. & Sorek, R. Widespread Utilization of Peptide Communication in Phages Infecting Soil and Pathogenic Bacteria. Cell Host and Microbe 25, 746–755.e5 (2019).

14. Gallego del Sol, F. Quiles-Puchalt, N., Brady, A., Penadés, J. R. & Marina, A. Insights into the mechanism of action of the arbitrium communication system in SPbeta phages. Nature communications 13, 1–15 (2022).

15. Doron, S. et al. Systematic discovery of antiphage defense systems in the microbial pangenome. Science 359, eaar4120 (2018).

16. Gao, L. et al. Diverse enzymatic activities mediate antiviral immunity in prokaryotes. Science 369, 1077–1084 (2020).

17. Millman, A. et al. An expanding arsenal of immune systems that protect bacteria from phages. bioRxiv (2022).

18. Tal, N. & Sorek, R. SnapShot: bacterial immunity. Cell 185, 578–578. e1 (2022).

19. Vassallo, C., Doering, C., Littlehale, M. L., Teodoro, G. & Laub, M. T. Mapping the landscape of anti-phage defense mechanisms in the E. coli pangenome. 2022.05.12.491691 Preprint at https://doi.org/10.1101/2022.05.12.491691 (2022).

20. Fillol-Salom, A. et al. Bacteriophages benefit from mobilizing pathogenicity islands encoding immune systems against competitors. Cell 185, 3248–3262.e20 (2022).

21. Lopatina, A., Tal, N. & Sorek, R. Abortive infection: bacterial suicide as an antiviral immune strategy. Annual review of virology 7, 371–384 (2020).

22. Pecota, D. C. & Wood, T. K. Exclusion of T4 phage by the hok/sok killer locus from plasmid R1. Journal of bacteriology 178, 2044–2050 (1996).

23. Song, S. & Wood, T. K. A primary physiological role of toxin/antitoxin systems is phage inhibition. Frontiers in Microbiology 11, 1895 (2020).

24. Jurėnas, D., Fraikin, N., Goormaghtigh, F. & Van Melderen, L. Biology and evolution of bacterial toxin–antitoxin systems. Nature Reviews Microbiology 20, 335–350 (2022).

25. LeRoux, M. & Laub, M. T. Toxin-Antitoxin Systems as Phage Defense Elements. Annual Review of Microbiology 76, p(2022).

26. Koga, M., Otsuka, Y., Lemire, S. & Yonesaki, T. Escherichia coli rnlA and rnlB Compose a Novel Toxin–Antitoxin System. Genetics 187, 123–130 (2011).

27. Fineran, P. C. et al. The phage abortive infection system, ToxIN, functions as a protein– RNA toxin–antitoxin pair. Proceedings of the National Academy of Sciences 106, 894– 899 (2009).

28. Dy, R. L., Przybilski, R., Semeijn, K., Salmond, G. P. & Fineran, P. C. A widespread bacteriophage abortive infection system functions through a Type IV toxin–antitoxin mechanism. Nucleic acids research 42, 4590–4605 (2014).

29. Tock, M. R. & Dryden, D. T. The biology of restriction and anti-restriction. Current opinion in microbiology 8, 466–472 (2005).

30. Hampton, H. G., Watson, B. N. & Fineran, P. C. The arms race between bacteria and their phage foes. Nature 577, 327–336 (2020).

31. Pinilla-Redondo, R. et al. Discovery of multiple anti-CRISPRs highlights anti-defense gene clustering in mobile genetic elements. Nature communications 11, 1–11 (2020).

32. Hobbs, S. J. et al. Phage anti-CBASS and anti-Pycsar nucleases subvert bacterial immunity. Nature 605, 522–526 (2022).

33. Ho, P., Chen, Y., Biswas, S., Canfield, E. & Feldman, D. E. Bacteriophage anti-defense genes that neutralize TIR and STING immune responses. bioRxiv (2022).

34. Otsuka, Y. & Yonesaki, T. Dmd of bacteriophage T4 functions as an antitoxin against Escherichia coli LsoA and RnlA toxins. Molecular microbiology 83, 669–681 (2012).

35. LeRoux, M. et al. The DarTG toxin-antitoxin system provides phage defence by ADP-ribosylating viral DNA. Nat Microbiol 7, 1028–1040 (2022).

36. Srikant, S., Guegler, C. K. & Laub, M. T. The evolution of a counter-defense mechanism in a virus constrains its host range. bioRxiv (2022).

37. Goldberg, G. W., Jiang, W., Bikard, D. & Marraffini, L. A. Conditional tolerance of temperate phages via transcription-dependent CRISPR-Cas targeting. Nature 514, 633– 637 (2014).

38. Rollie, C. et al. Targeting of temperate phages drives loss of type I CRISPR–Cas systems. Nature 578, 149–153 (2020).

39. Varble, A. et al. Prophage integration into CRISPR loci enables evasion of antiviral immunity in Streptococcus pyogenes. Nat Microbiol 6, 1516–1525 (2021).

40. Simanshu, D. K., Yamaguchi, Y., Park, J.-H., Inouye, M. & Patel, D. J. Structural basis of mRNA recognition and cleavage by toxin MazF and its regulation by antitoxin MazE in Bacillus subtilis. Molecular cell 52, 447–458 (2013).

41. Kohm, K. et al. The Bacillus phage SPβ and its relatives: A temperate phage model system reveals new strains, species, prophage integration loci, conserved proteins and lysogeny management components. Environmental Microbiology 24, 2098–2118 (2022).

42. Nicolas, P. et al. Condition-Dependent Transcriptome Reveals High-Level Regulatory Architecture in Bacillus subtilis. Science 335, 1103–1106 (2012).

43. Zhu, B. &Stülke, J. SubtiWiki in 2018: from genes and proteins to functional network annotation of the model organism Bacillus subtilis. Nucleic Acids Res 46, D743–D748 (2018).

44. Pellegrini, O., Mathy, N., Gogos, A., Shapiro, L. & Condon, C. The Bacillus subtilis ydcDE operon encodes an endoribonuclease of the MazF/PemK family and its inhibitor. Molecular microbiology 56, 1139–1148 (2005).

45. Park, J.-H., Yamaguchi, Y. & Inouye, M. Bacillus subtilis MazF-bs (EndoA) is a UACAU-specific mRNA interferase. FEBS letters 585, 2526–2532 (2011).

46. Johnson, C. M., Harden, M. M. & Grossman, A. D. Interactions between mobile genetic elements: An anti-phage gene in an integrative and conjugative element protects host cells from predation by a temperate bacteriophage. PLoS Genetics 18, e1010065 (2022).

47. Hazan, R. & Engelberg-Kulka, H. Escherichia coli mazEF-mediated cell death as a defense mechanism that inhibits the spread of phage P1. Molecular Genetics and Genomics 272, 227–234 (2004).

48. Alawneh, A. M., Qi, D., Yonesaki, T. & Otsuka, Y. An ADP-ribosyltransferase Alt of bacteriophage T4 negatively regulates the Escherichia coli MazF toxin of a toxin– antitoxin module. Molecular Microbiology 99, 188–198 (2016).

49. Guegler, C. K. & Laub, M. T. Shutoff of host transcription triggers a toxin-antitoxin system to cleave phage RNA and abort infection. Molecular Cell 81, 2361–2373. e9 (2021).

50. Bobonis, J. et al. Bacterial retrons encode phage-defending tripartite toxin–antitoxin systems. Nature 1–3 (2022).

51. Wu, X., Wang, X., Drlica, K. & Zhao, X. A Toxin-Antitoxin Module in Bacillus subtilis Can Both Mitigate and Amplify Effects of Lethal Stress. PLOS ONE 6, e23909 (2011).

52. Hochhauser, D., Millman, A. & Sorek, R. The defence island repertoire of the Escherichia coli pan-genome. 2022.06.09.495481 Preprint at https://doi.org/10.1101/2022.06.09.495481 (2022).

53. Brito, P. H. et al. Genetic Competence Drives Genome Diversity in Bacillus subtilis. Genome Biology and Evolution 10, 108–124 (2018).

54. Harwood, C. R. & Cutting, S. M. Molecular biological methods for Bacillus. (Wiley, 1990).

55. McDonnell, G. E., Wood, H., Devine, K. M. & McConnell, D. J. Genetic control of bacterial suicide: regulation of the induction of PBSX in Bacillus subtilis. J Bacteriol 176, 5820–5830 (1994).

56. Koo, B.-M. et al. Construction and Analysis of Two Genome-Scale Deletion Libraries for Bacillus subtilis. Cell Systems 4, 291–305.e7 (2017).

57. Gibson, D. G. et al. Enzymatic assembly of DNA molecules up to several hundred kilobases. Nat Methods 6, 343–345 (2009).

58. Webb, B. & Sali, A. Comparative protein structure modeling using MODELLER. Current protocols in bioinformatics 54, 5.6. 1-5.6. 37 (2016).

59. Glaser, F. et al. ConSurf: identification of functional regions in proteins by surfacemapping of phylogenetic information. Bioinformatics 19, 163–164 (2003).

60. Ashkenazy, H. et al. ConSurf 2016: an improved methodology to estimate and visualize evolutionary conservation in macromolecules. Nucleic acids research 44, W344–W350 (2016).

61. Ko, J., Park, H., Heo, L. & Seok, C. GalaxyWEB server for protein structure prediction and refinement. Nucleic acids research 40, W294–W297 (2012).

