## Supplementary data for "Arbitrium communication controls phage life-cycle through modulation of a bacterial anti-phage defense system"

### Supplementary Figures

```
aimXORF     1   atgaaaagagcattaggtaaagcaatatcttatgaagaaatggcaaaagggtacgaggaa 60
aimXrecoded 1   atgaagcgcgcgttgggaaaggcgatctcatacgaagagatggcgaagggatatgaagag 60
                 M  K  R  A  L  G  K  A  I  S  Y  E  E  M  A  K  G  Y  E  E

aimXORF     61  atggctgcaatcaattcaataattgctcaagaggacaaccatcttgagaatgaagcggaa 120
aimXrecoded 61  atggcagctattaactctatcattgcacaggaagataatcatttagaaaacgaggcagag 120
                 M  A  A  I  N  S  I  I  A  Q  E  D  N  H  L  E  N  E  A  E

aimXORF     121 atgattaaaacaaggtataaaaccctggcttcatga 156
aimXrecoded 121 atgatcaaaacgagatacaaaacacttgcatcttga 156
                 M  I  K  T  R  Y  K  T  L  A  S  *
```

**Extended Data Figure 1. *aimX*$^{ORF}$ and *aimX*$^{recoded}$.** DNA sequence of *aimX*$^{ORF}$ and of *aimX*$^{recoded}$ together with the amino acid sequence.

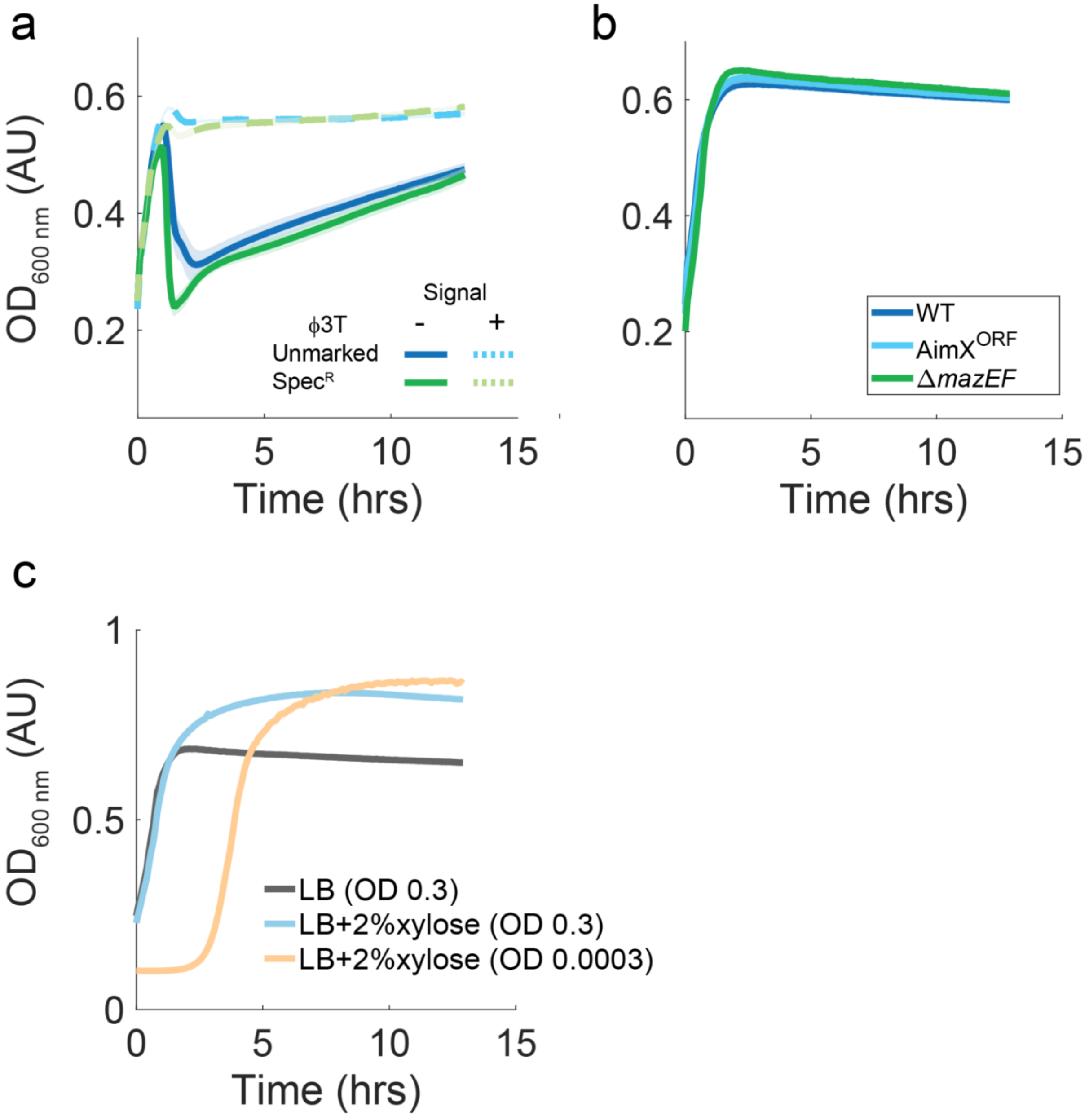


**Extended Data Figure 2. Control growth curves and plaque counts.** (A) growth curve of the wild-type bacterial strain infected with either unmarked ϕ3T (blue), or a spectinomycin marked ϕ3T (green) either with (dashed) or without (solid line) the addition of the arbitrium signal. The phages behave similarly in their growth patterns and response to the arbitrium signal. (B) Growth of strains in the absence of infection. Shown are wild-type, AimXORF with 100µM IPTG, Δ*mazEF*, Δ*mazF* . (C) growth of the wild-type strain with or without addition of 2% xylose to LB medium. Growth was initiated at OD 0.3 or after a 1000-fold dilution of OD 0.3 as marked in the legend. All growth experiments were done on multiple biological repeats. Shaded area marks the biological standard error of these repeats where in each day the curve is an average over multiple technical repeats.

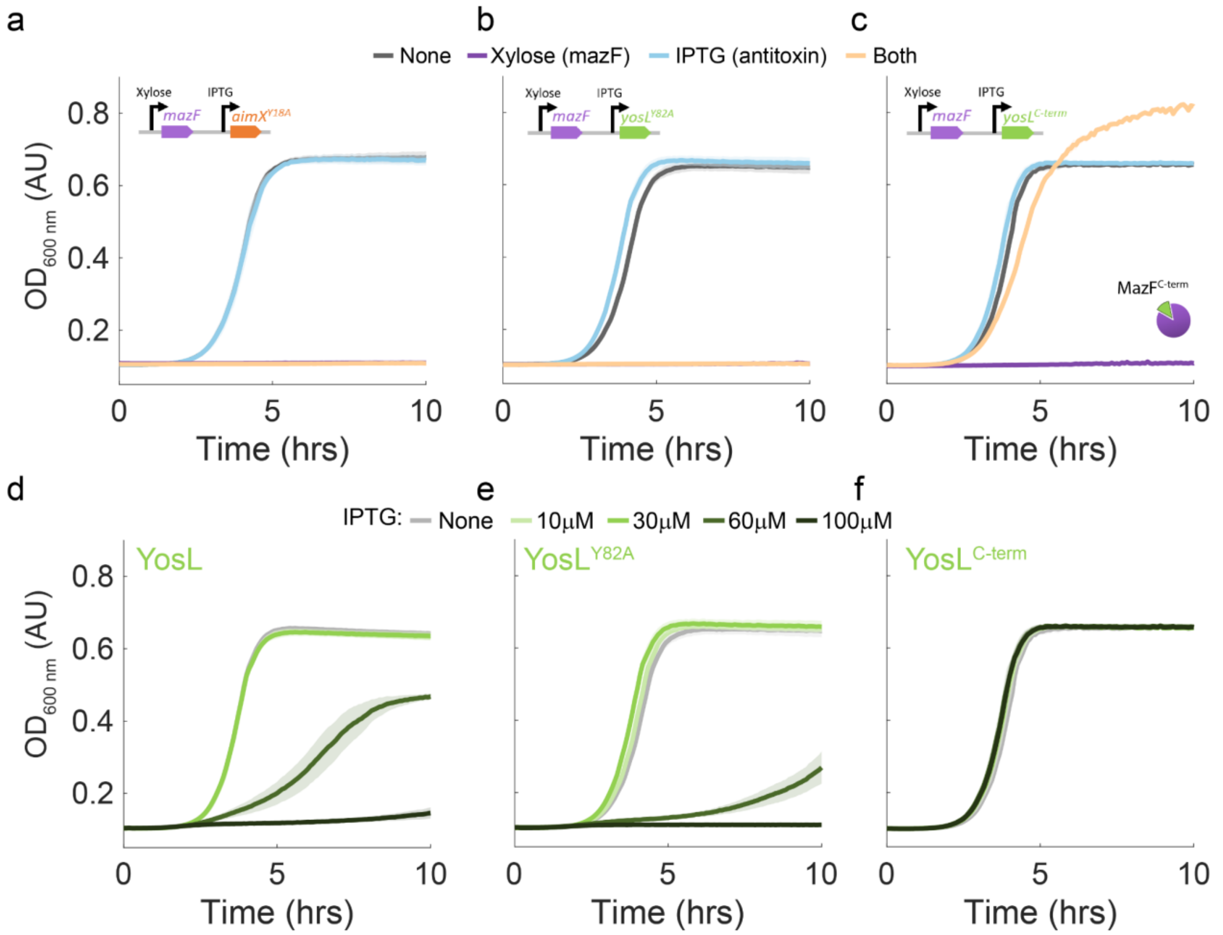


**Extended Data Figure 3: Mutations and manipulation of expression of *aimX* and *yosL*.** (A-C) expression of antitoxin mutants with or without MazF. Growth curves of cells with addition of 100µM IPTG (Antitoxin, light blue), 2% xylose (MazF, purple), none (gray) or both (light orange) for AimX (A), YosL (B) (As in main figure 1 panels F,G), but with the tyrosine in position 18 of AimX and position ,82 of YosL mutated to alanine. (C) shows expression of the C-terminal part of YosL. Note that the xylose only (purple) and xylose and IPTG (light orange) overlap in panels (A,B). (C,D) Growth curves of YosL (D) $YosL^{Y82A}$ (E) and $YosL^{C\text{-}term}$ (F) as a function of time with different levels of IPTG, as indicated in the legend. All growth experiments were done on multiple biological repeats. Shaded area marks the biological standard error of these repeats where in each day the curve is an average over multiple technical repeats.

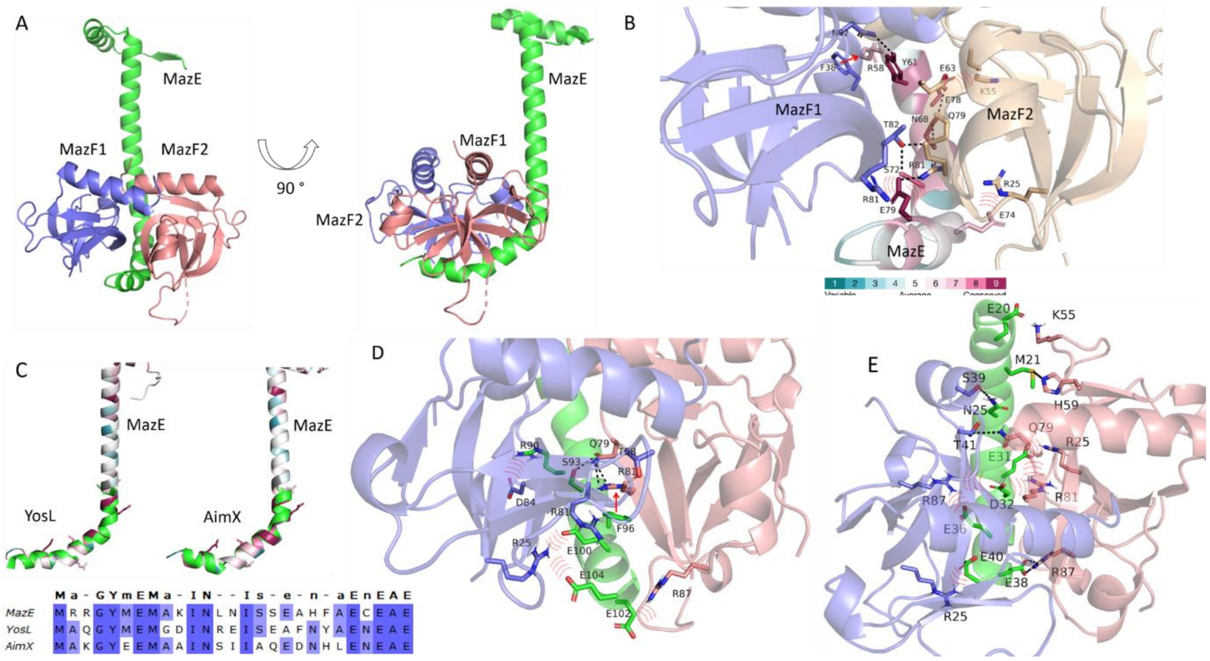


**Extended Data Figure 4: Homology modeling of AimX and YosL binding to a MazF dimer.** Global (A) and detailed (B) structure of MazE and a mazF dimer, derived from the crystal structure of the proteins (PDB ID: 4me7)[40]. In (A), the global structure is shown from two different angles, where the three chains are shown as ribbons, with MazE colored green and the two MazF chains colored blue and light pink. In (B), MazE is colored by evolutionary conservation (see color code at the bottom of the image), calculated using ConSurf[59], and MazF is shown as in (A). The polar side chain interactions between MazE and MazF are noted as follows: hydrogen bonds are shown as black dashed lines, ionic interactions are shown as red curves, and the cation-π interaction is shown as a red arrow. (C-E) global (C) and detailed (D,E) predicted structures of YosL (C-left, D) and AimX (C-right, E). The structures were predicted by homology modeling, based on the crystal structure of the MazE-MazF$_2$ dimer (Methods). The sequence homology between MazE and the two phage proteins, on which the modeling relied, is also shown (C-bottom). No considerable changes to the structure are observed for YosL and AimX and many of the interactions are preserved. Amino-acid with specific interactions between antitoxin and MazF are shown in (D-F).

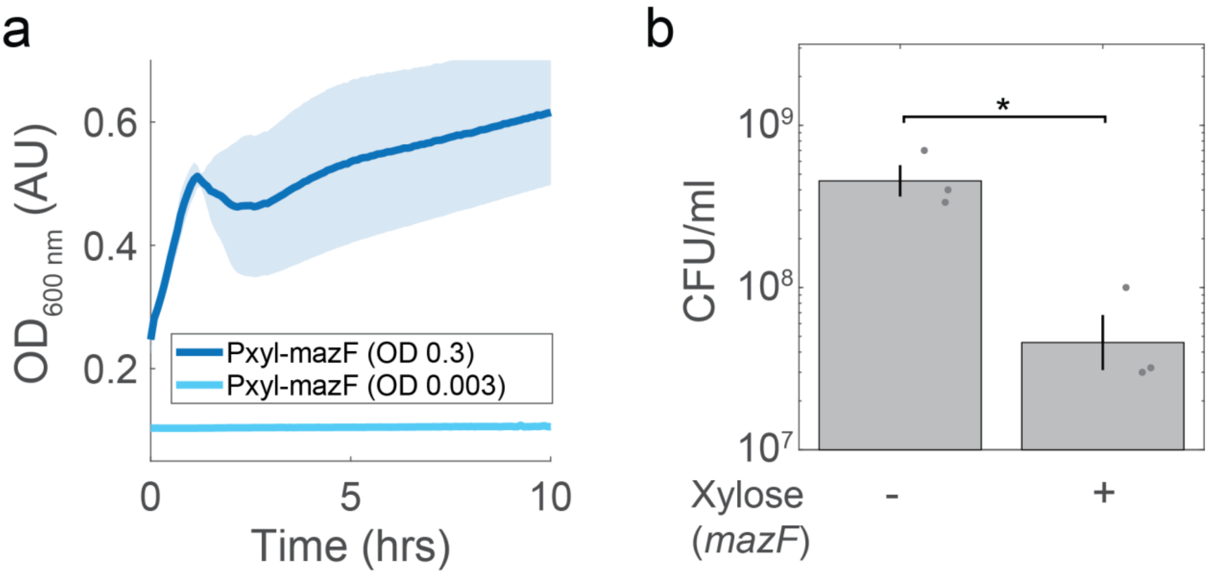


**Extended Data Figure 5: MazF induction at mid-log phase does not lead to immediate cessation of growth.** (A) growth curve of $P_{xyl}$-*mazF* cells induced at low OD (light blue, taken from Fig. 1F) and at an OD=0.3 (dark blue) with 2% xylose. (B) $P_{xyl}$-*mazF* cells were either added or not added 2% xylose at an OD of 0.3 for two hours and then plated on plates without xylose. Colony forming units were counted after an overnight growth. CFU with addition of xylose is smaller by an order of magnitude (p=0.007, t-test, N=3). All growth experiments were done on multiple biological repeats. Shaded area marks the biological standard error of these repeats where in each day the curve is an average over multiple technical repeats.

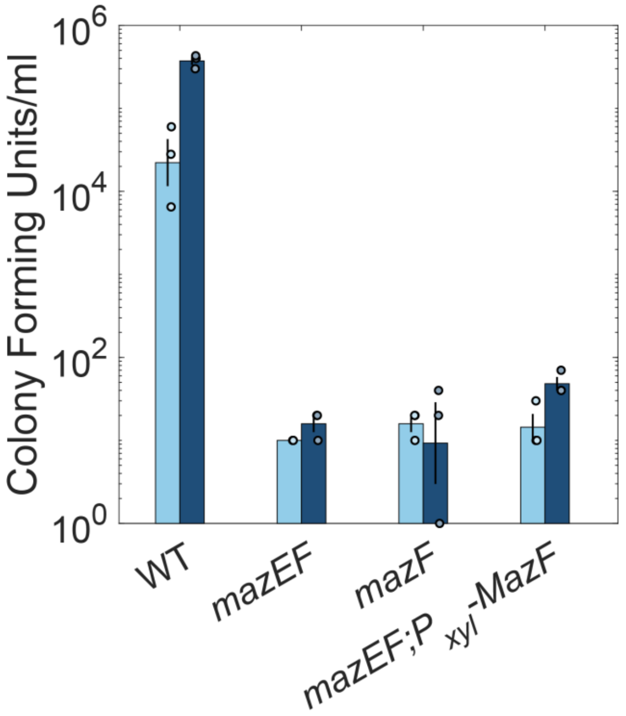


**Extended Data Figure 6: Lysogens count 20 minutes after infection.** As in main figure 2D, but 20 minutes after infection instead of two hours.

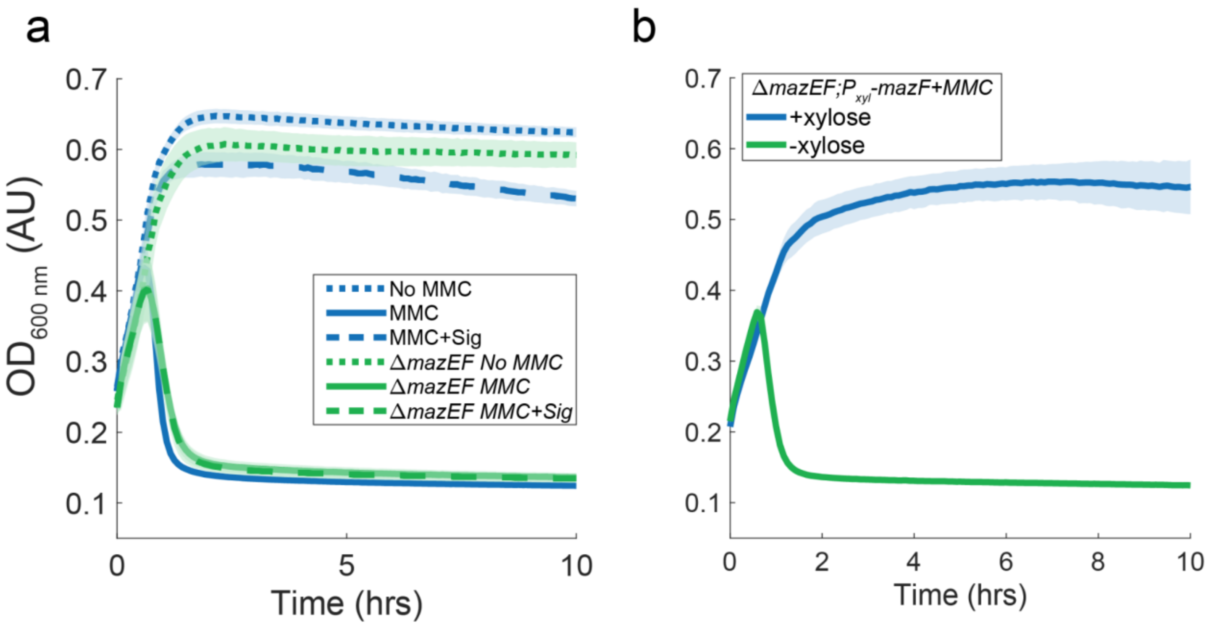


**Extended Data Figure 7: MazEF affects prophage induction in addition to infection.** (A) Induction curves for two lysogenic strains; wild-type (blue), Δ*mazEF* (green). Mitomycin C (MMC) was added at time zero at a concentration of 500µg/ml either without (solid line) or with (dashed line) the arbitrium peptide (SAIRGA). Also shown are growth curves with no prophage induction (dotted) (B) Induction curves for a Δ*mazEF;P*$_{xyl}$-*mazF* with (blue) or without (green) the addition of xylose. MMC was added at time 0 at a concentration of 500µg/ml. Xylose was added at the same time as MMC at 2%. All growth experiments were done on multiple biological repeats. Shaded area marks the biological standard error of these repeats where in each day the curve is an average over multiple technical repeats.

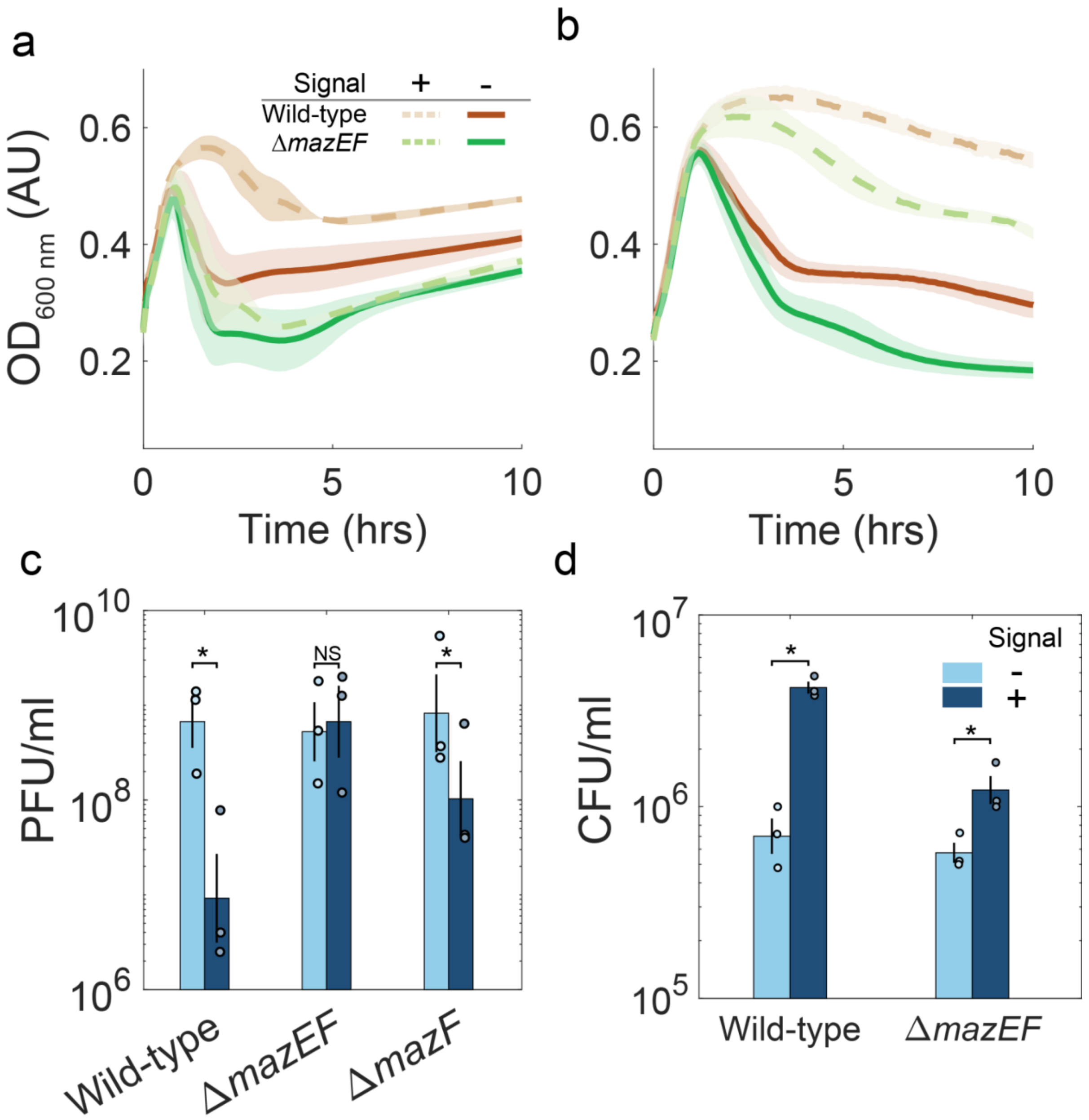


**Extended Data Figure 8: Impact of the mazEF system of phage SPβ lysis/lysogeny.** (A,B) Growth curve during infection (A) or prophage induction (B) of wild-type (brown) and Δ*mazEF* (green) cells. In (A) cells were infected at time 0 with phage SPβ. In (B) SPβ lysogens were induced at time 0 with 500µg/ml MMC. Both were done either with (dashed line) or without (solid line) the SPβ arbitrium peptide (GMPRGA). (C) Plaque counts after overnight SPβ infection of wild-type, Δ*mazEF or* Δ*mazF* with an arbitrium signal either added (dark blue) or not added (light blue) at time of infection. (D) Lysogen numbers 20 minutes after infection with marked SPβ of wild-type or ΔmazEF cells with an arbitrium signal either added (dark blue) or not added (light blue) at time of infection.

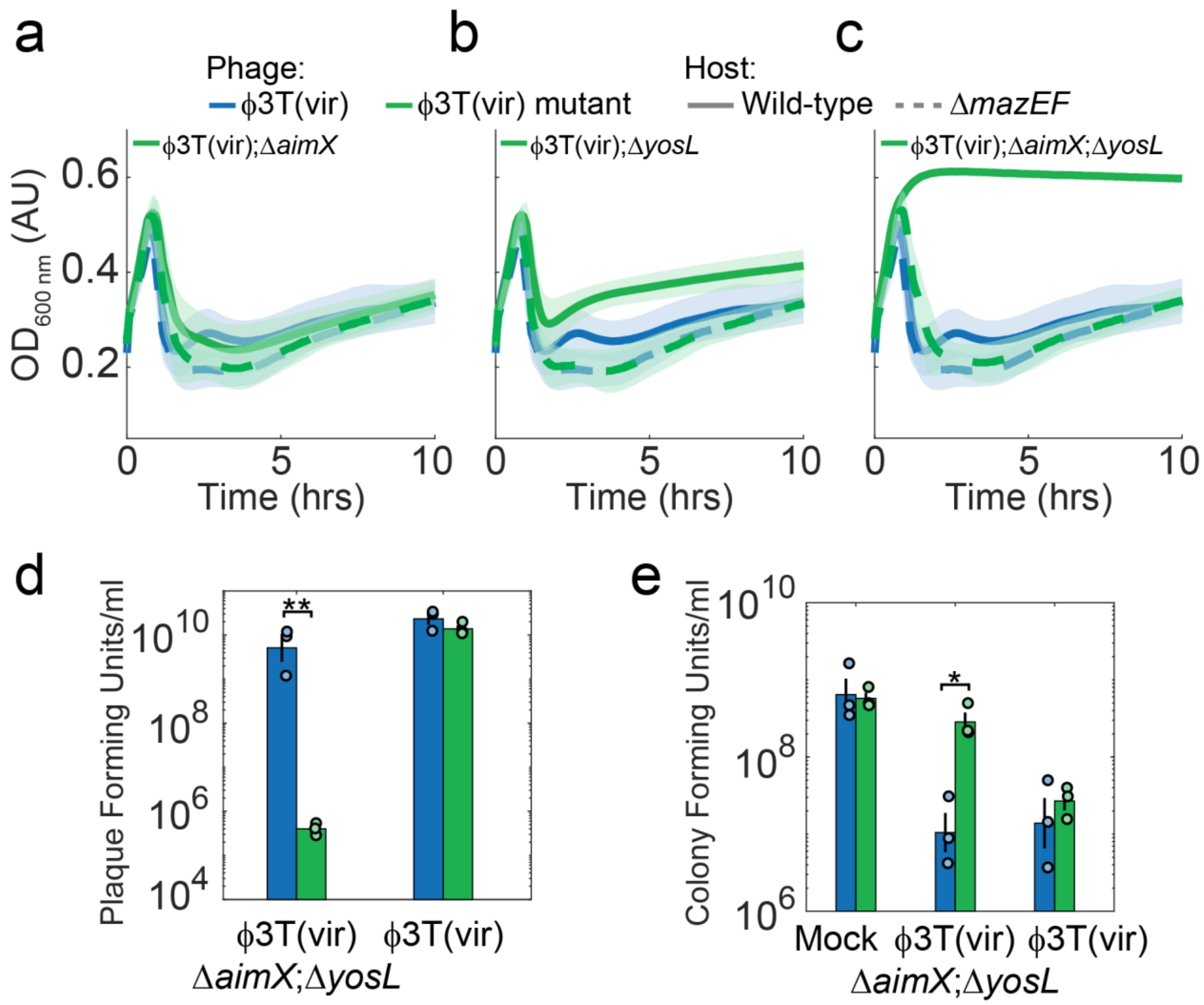


**Extended Data Figure 9: MazEF prevents infection of a lytic ϕ3T variant lacking antitoxins.** (A-C) Infection curves of strain ϕ3T(vir) (blue) compared with mutant strains (green) (A) ϕ3T(vir);Δ*aimX*, (B) ϕ3T(vir);Δ*yosL*, (C) ϕ3T(vir);Δ*aimX;*Δ*yosL*. The ϕ3T(vir) graph is the same for all panels. Each of the phage strains infected either wild-type (solid) or *ΔmazEF* (dashed) bacterial hosts. (D,E) Plaque (D) and colony (E) counts 2 hours after infection of a ϕ3T(vir);Δ*aimX;*Δ*yosL* and ϕ3T(vir) strains at a MOI of 4 on wild-type (green) and *ΔmazEF* (blue) bacteria. Colony forming units were also measured for a mock infection (marked Mock).

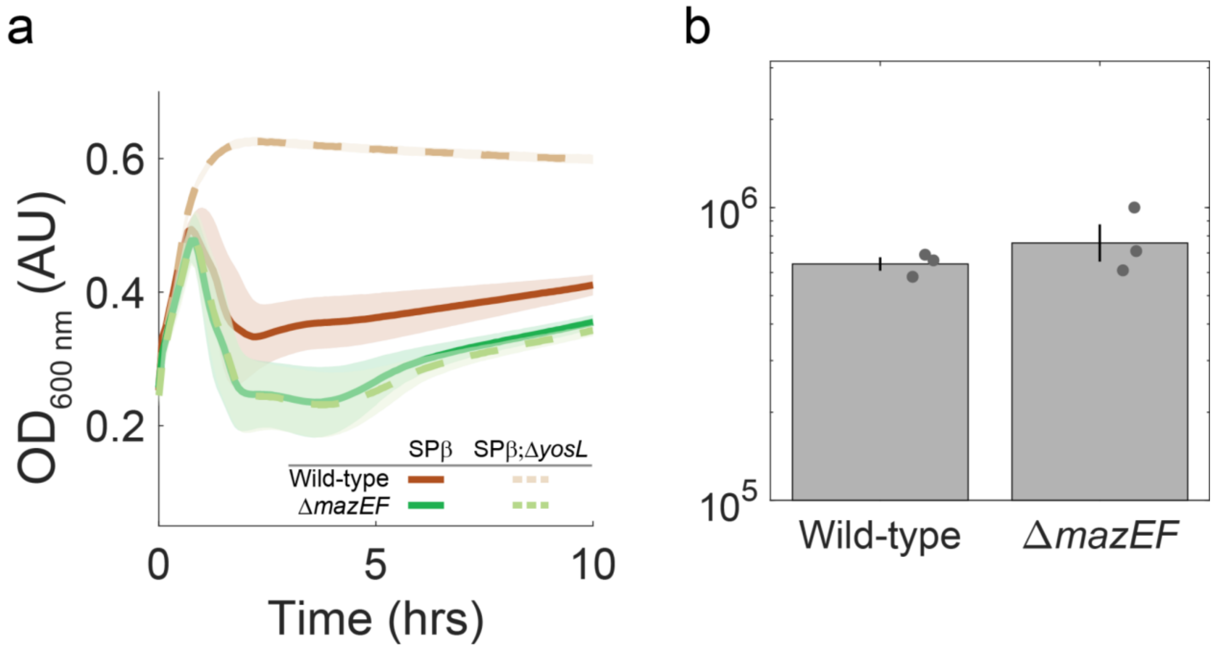


**Extended Data Figure 10: MazEF defends against a marked phage SPβ;*yosL* mutant.** (A) Infection at MOI 0.1 of wild-type (brown) and Δ*mazEF* (green) cells with either wild-type phage SPβ (solid line) or a SPβ;Δ*yosL* mutant (dashed line). (B) Lysogen numbers during infection of spectinomycin resistance marked SPβ;Δ*yosL* (ref [46]) Infecting either wild-type or Δ*mazEF* mutant cells. Infected cells were plated on a spectinomycin plate 2 hours after infection.

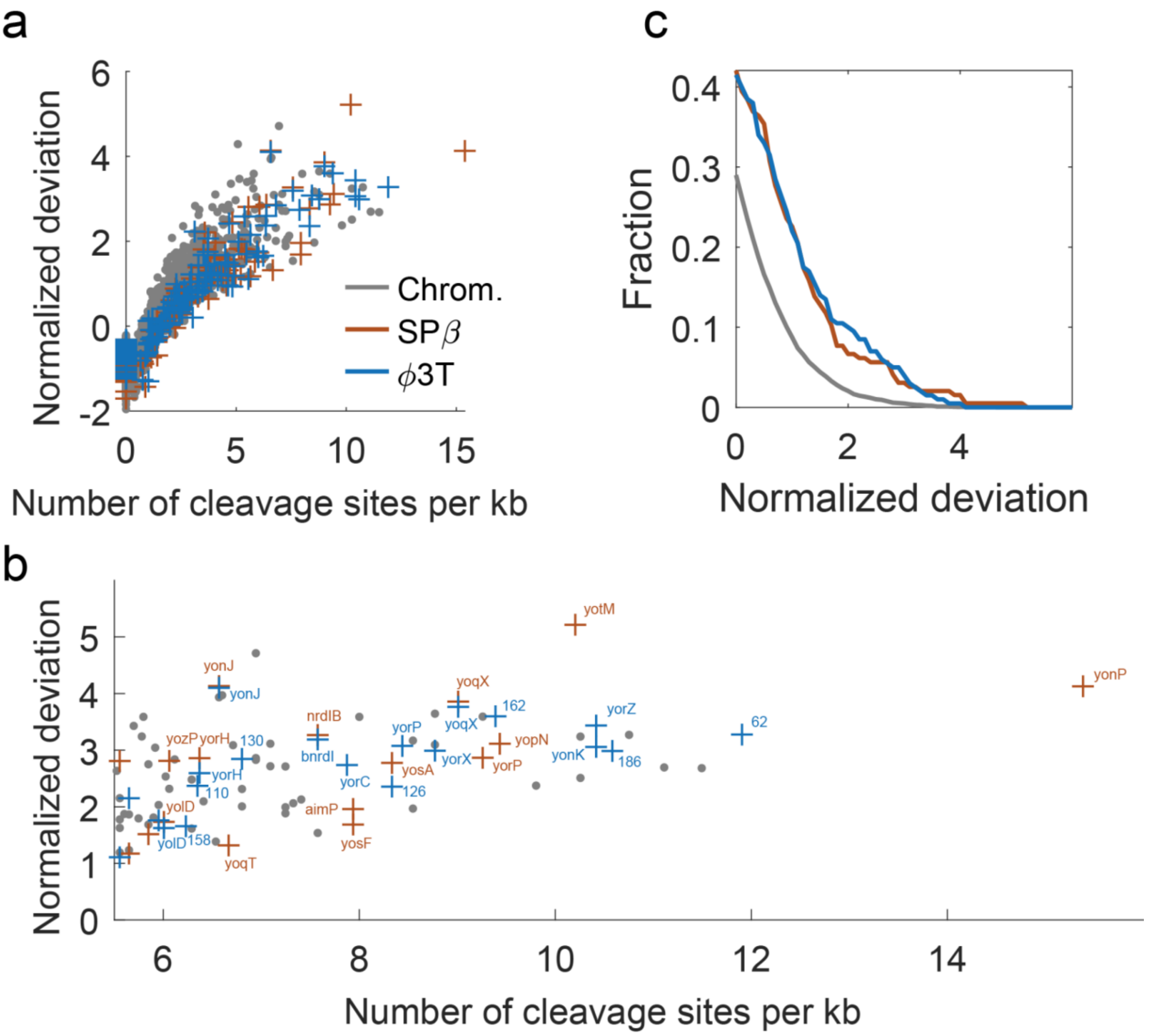


**Extended Data Figure 11: MazF cleavage site statistics on *B. subtilis* chromosome and phages SPβ and ϕ3T.** (A,B) Each chromosomal gene (gray dot), or phage SPβ (brown) and ϕ3T (blue) gene is plotted on an x-y axis showing the number of MazF cleavage sites per kb (UACAU) on the open reading frame of the gene on the x-axis. The y-axis marks the normalized deviation of the number of cleavage sites on the gene. For each gene we randomly reshuffled the ORF sequence a 1000 times and calculated mean and standard deviation. The value shown is the number of cleavage sites in the real gene minus the average number in the reshuffled collection divided by the standard deviation of this number. In (A) all genes are shown. In (B) only genes whose x-axis value is above 5.5 are shown. All SPβ and ϕ3T genes above a value of 6 are named. SPβ gene names are based on their naming in *B. subtilis* str. 168 (accession NC_000964.3). ϕ3T gene names are based on the ncbi sequence (accession KY030782). If a gene is annotated as homologous to a SPβ gene, its homolog's name is given, otherwise, its number is given.
